# Light-triggered protease-mediated release of actin-bound cargo from synthetic cells

**DOI:** 10.1101/2024.09.15.613133

**Authors:** Mousumi Akter, Hossein Moghimianavval, Gary D. Luker, Allen P. Liu

**Affiliations:** Department of Mechanical Engineering, University of Michigan, Ann Arbor, MI, 48109, USA; Department of Biomedical Engineering, University of Michigan, Ann Arbor, MI, 48109, USA; Department of Radiology, University of Michigan, Ann Arbor, MI, 48109, USA; Cellular and Molecular Biology Program, University of Michigan, Ann Arbor, MI, 48109, USA; Department of Biophysics, University of Michigan, Ann Arbor, MI, 48109, USA

## Abstract

Synthetic cells offer a versatile platform for addressing biomedical and environmental challenges, due to their modular design and capability to mimic cellular processes such as biosensing, intercellular communication, and metabolism. Constructing synthetic cells capable of stimuli-responsive secretion is vital for applications in targeted drug delivery and biosensor development. Previous attempts at engineering secretion for synthetic cells have been confined to non-specific cargo release via membrane pores, limiting the spatiotemporal precision and specificity necessary for selective secretion. Here, we designed and constructed a protein-based platform termed TEV Protease-mediated Releasable Actin-binding protein (TRAP) for selective, rapid, and triggerable secretion in synthetic cells. TRAP is designed to bind tightly to reconstituted actin networks and is proteolytically released from bound actin, followed by secretion via cell-penetrating peptide membrane translocation. We demonstrated TRAP’s efficacy in facilitating light-activated secretion of both fluorescent and luminescent proteins. By equipping synthetic cells with a controlled secretion mechanism, TRAP paves the way for the development of stimuli-responsive biomaterials, versatile synthetic cell-based biosensing systems, and therapeutic applications through the integration of synthetic cells with living cells for targeted delivery of protein therapeutics.

## 1 Introduction

Synthetic cells are non-living cell mimics designed and engineered to tackle modern biomedical and environmental challenges.^[1–3]^ The modular nature of synthetic cells and the bottom-up approach of their synthesis enable the inclusion of different processes, ranging from biosensing^[4,5]^ to intercellular communication,^[6–8]^ that collectively define the function and specialization of the synthetic cell.^[9]^ The biomimicry of metabolism,^[10]^ energy production,^[11]^ cytoskeleton,^[12]^ and cell-free protein synthesis^[13]^ in synthetic cells has paved the way for generating sophisticated synthetic cells aimed at engineering and basic science applications.

Nevertheless, the construction of synthetic cells for creating stimuli-responsive biomaterials, triggerable release of molecules, or building intricate biosensors could only be realized if synthetic cells can sense, process, and respond to external stimuli. Specifically, the ability to secrete molecules in response to inputs unlocks novel forms of intercellular communication, targeted delivery of protein biologics, and stimuli-responsive biosensors. Secretory synthetic cells may also find use in industrial applications, such as high throughput cell-free synthesis of proteins and antibodies.

Given its vital role in numerous biological processes^[14–23]^, it is not surprising that cellular secretion stands at the center of many studies in top-down synthetic biology. Recombinant secretory expression of insulin in *Saccharomyces cerevisiae* accounts for producing half of the global insulin supply.^[24]^ Top-down engineering of cellular secretory pathways has enabled programmable cellular secretion with therapeutic applications.^[25,26]^ However, compared to top-down systems, biomimicry of secretion in bottom-up synthetic cells has been mostly limited to non-specific release of molecules through membrane pores.^[27–34]^ This is perhaps due to the fact that imitating the exact mechanism of cellular secretion, which relies on vesicle fusion, is not trivial.

Pore-forming toxins such as α-hemolysin or perfringolysin O, as well as membrane active peptides like melittin, have been the main tools for enabling synthetic cells to secrete molecules.^[35–37]^ Release of small molecules such as isopropyl β-d-1-thiogalactopyranoside (IPTG) or proteins like brain-derived neurotrophic factor from synthetic cells through α-hemolysin^[35]^ and perfringolysin O,^[36]^ respectively, has been utilized to demonstrate communication between synthetic cells and bacteria or neurons. Similarly, the addition of melittin to synthetic cells is widely used to trigger secretion, which initiates enzymatic reaction cascades or induces signaling among synthetic cells.^[37]^

While cargo release through pores is effective and can be genetically programmed (*i*.*e*., through cell-free expression of the pore subunits), secretion *via* this approach is limited only to molecules smaller than the pore size. Additionally, pores made by toxins and membrane-active peptides allow the translocation of molecules across the bilayer non-selectively and potentially could damage adjacent living cells. The release of internal molecules through membrane rupture^[38,39]^ shares this limitation with membrane pores as well.

Natural cells cleverly regulate selective secretion by using specific protein-protein interactions that drive fusion between cargo-carrying vesicles and specific target organelles or membranes.^[40]^ Additionally, the inclusion of secretion signal peptides on the nascent protein chain ensures the translocation of secretory proteins to the endoplasmic reticulum, priming them for release to the extracellular environment.^[41]^ While a few studies have implemented vesicle fusion to demonstrate secretion from synthetic cells^[42]^ or synthetic endocytosis,^[43,44]^ the complexity of the natural secretory pathway makes imitation of selective secretion based on vesicle fusion challenging.

It has been demonstrated that membrane translocation domain of cell-free synthesized exotoxin A of *Pseudomonas aeruginosa* can mediate pore-free secretion from synthetic cells.^[45]^ A recent work by Heili *et al*. utilized a similar approach by fusing a cell-penetrating peptide (CPP) domain to the cell-free expressed cargo encapsulated in synthetic cells, thus enabling CPP-mediated membrane translocation and secretion of cargos as large as 100 kDa.^[46]^ Notably, the CPP tag penetratin, used by Heili *et al*., is not a pore-forming CPP.^[47–49]^ It translocates fusion proteins across the membrane without causing non-selective leakage of synthetic cell contents. Addition of tags like CPP to the cargo to enable secretion promises a simple, yet effective approach to encode selective, leakless secretion. However, secretion *via* cell-free synthesis of cargo is slow as the transcription and translation typically take a few hours. Secretion via cell-free synthesis also lacks temporal control of cargo release.

Here, we introduce a TEV protease-mediated protein for rapid, triggerable selective secretion of molecules from synthetic cells. We utilize CPP-mediated membrane translocation to enable selective cargo secretion. To make the initiation of secretion triggered by external stimuli, we designed the cargo such that it is tightly bound to a large, soluble scaffold inside the synthetic cell. Any stimuli that cause unbinding of the cargo from the scaffold (*e*.*g*., through protease activity) would induce cargo secretion from the synthetic cell. While we demonstrate light-activated cargo secretion here, the platform allows modular design of cargos, thus enabling multiplexed cargo secretion in response to combinatorial inputs. We envision this platform to be a useful tool for intercellular communication between synthetic cells as well as therapeutic applications such as synthetic cell-natural cell communication or targeted release of biologically relevant peptides and proteins.

## 2 Results

### 2.1 TEV Protease-mediated Releasable Actin-binding Protein (TRAP) design

To design a secretion mechanism for synthetic cells, we leveraged the unique capability of CPP in translocating proteins across a lipid bilayer. CPPs are short natural or synthetic peptides that can cross the lipid bilayer. Naturally, CPPs are found in proteins such as toxins or viral proteins, facilitating the invasion or internalization of pathogenic proteins inside cells^[50]^. Among different CPPs, we chose penetratin because of its shown high translocation efficiency for cargos as large as 100 kDa in a pore-independent manner.^[46–49,51]^ Thus, we hypothesized that fusing penetratin to a cargo of interest will enable leakless cargo secretion from synthetic cells, as others have recently shown.^[46]^

Since CPP will passively allow diffusion of the cargo from synthetic cells as soon as encapsulated, we reasoned that a mechanism to retain the cargo inside the synthetic cell is required for regulated secretion. We hypothesized that if the cargo is tightly bound to a large, soluble scaffold inside the synthetic cell, it would not be able to freely diffuse across the bilayer. In other words, a ‘cargo sponge’ inside the synthetic cells would sequester the cargo, retaining it inside the synthetic cell, until a stimulus drives detachment of the cargo from the scaffold. We selected actin networks as the scaffold and reasoned that fusion of a high-affinity actin-binding domain to the cargo would allow its tight binding to the scaffold. Lastly, we reasoned that protease cleavage of the actin-binding domain from the cargo would enable its detachment from the scaffold, thus initiating its secretion from the synthetic cell. For the proof-of-concept demonstration, we chose TEV protease (TEVp) as the proteolytic enzyme for cargo detachment from the actin scaffold. Lastly, we selected giant unilamellar vesicles (GUVs) as our model synthetic cells due to their cell-like size and lipid bilayer membrane.

Since our designed protein-based secretion platform relies on actin-based sequestration and TEVp-mediated detachment, we named our secretion system <u>T</u>EVp-mediated releasable <u>a</u>ctin-binding <u>p</u>rotein, referred to as TRAP hereafter. **Figure 1a** illustrates the TRAP construct, which comprises a cargo of interest flanked by two domains: an N-terminal CPP domain and a C-terminal Lifeact domain, responsible for anchoring TRAP to the actin. Lifeact is a 17-amino-acid peptide widely used for staining filamentous actin in eukaryotic cells.^[52]^ We conducted all experiments with fascin-bundled actin (F-actin) for enhanced visualization by fluorescence microscopy. The cargo domain is separated from the Lifeact domain by a TEVp cleavage site (TCS), which allows for precise proteolytic cleavage by TEVp. As a control, we designed TRAP-ΔCPP, a variant lacking the N-terminal CPP domain but retaining the cargo and the C-terminal TCS-Lifeact, rendering it non-releasable from the GUV.

**Figure 1.**
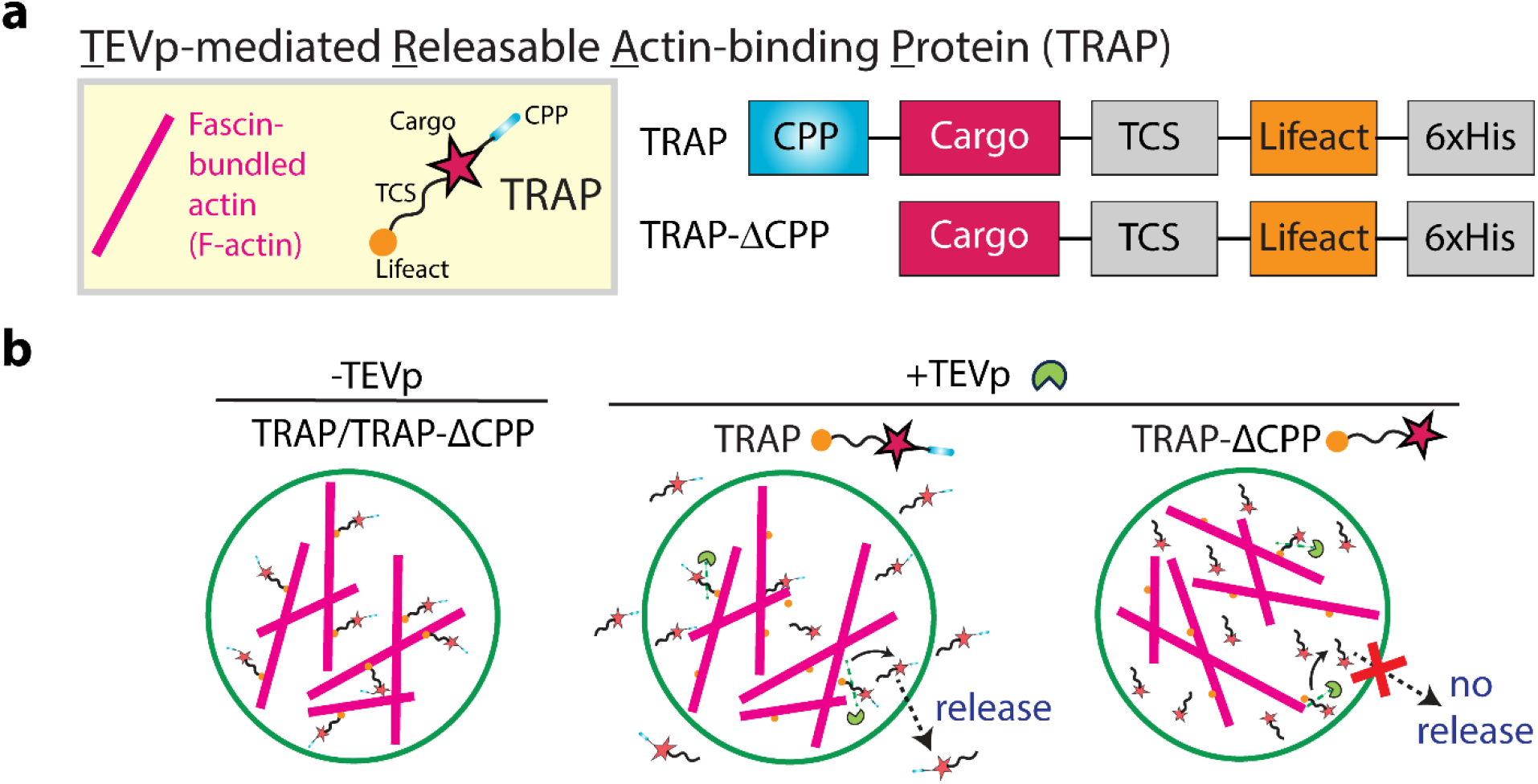
TEVp-mediated cargo release from GUVs. **a**. The left panel shows a schematic representation of fascin-bundled actin (F-actin) and TEVp-mediated releasable actin-binding protein (TRAP). The right panel illustrates the constructs for TRAP design: TRAP contains a cargo protein flanked by a cell-penetrating peptide (CPP) at the N-terminus and an actin-binding domain Lifeact at the C-terminus separated from the cargo by a TEVp cleavage site (TCS). TRAP-ΔCPP contains the cargo and the C-terminal TCS-actin-binding domain but lacks the N-terminal CPP domain. **b**. Schematic representation of cargo release from a GUV through unbinding of TRAP from the actin bundle *via* TEVp. TRAP is released from the GUV through its CPP domain while TRAP-ΔCPP remains inside the GUV.

**Figure 1b** shows the schematic representation of the mechanism underlying TEVp-mediated cargo release from GUVs. In the presence of TEVp, the TCS within TRAP is cleaved, leading to the detachment of the CPP-cargo from Lifeact, thereby releasing TRAP from F-actin. Subsequently, the CPP domain of TRAP facilitates its translocation across the GUV membrane, releasing the cargo into the extracellular environment. In contrast, TRAP-ΔCPP, lacking the CPP domain, remains confined within the GUV.

### 2.2 TRAP tightly binds to actin and can be detached from actin by TEVp

We first sought to test whether actin can sequester TRAP. As a model cargo protein, we chose mCherry as it allows for detecting TRAP by fluorescence microscopy. We created constructs TRAP-mCherry and its control variant TRAP-ΔCPP-mCherry (**Figure 2a**) and purified them from *Escherichia coli* (**Figure S1**).

**Figure 2.**
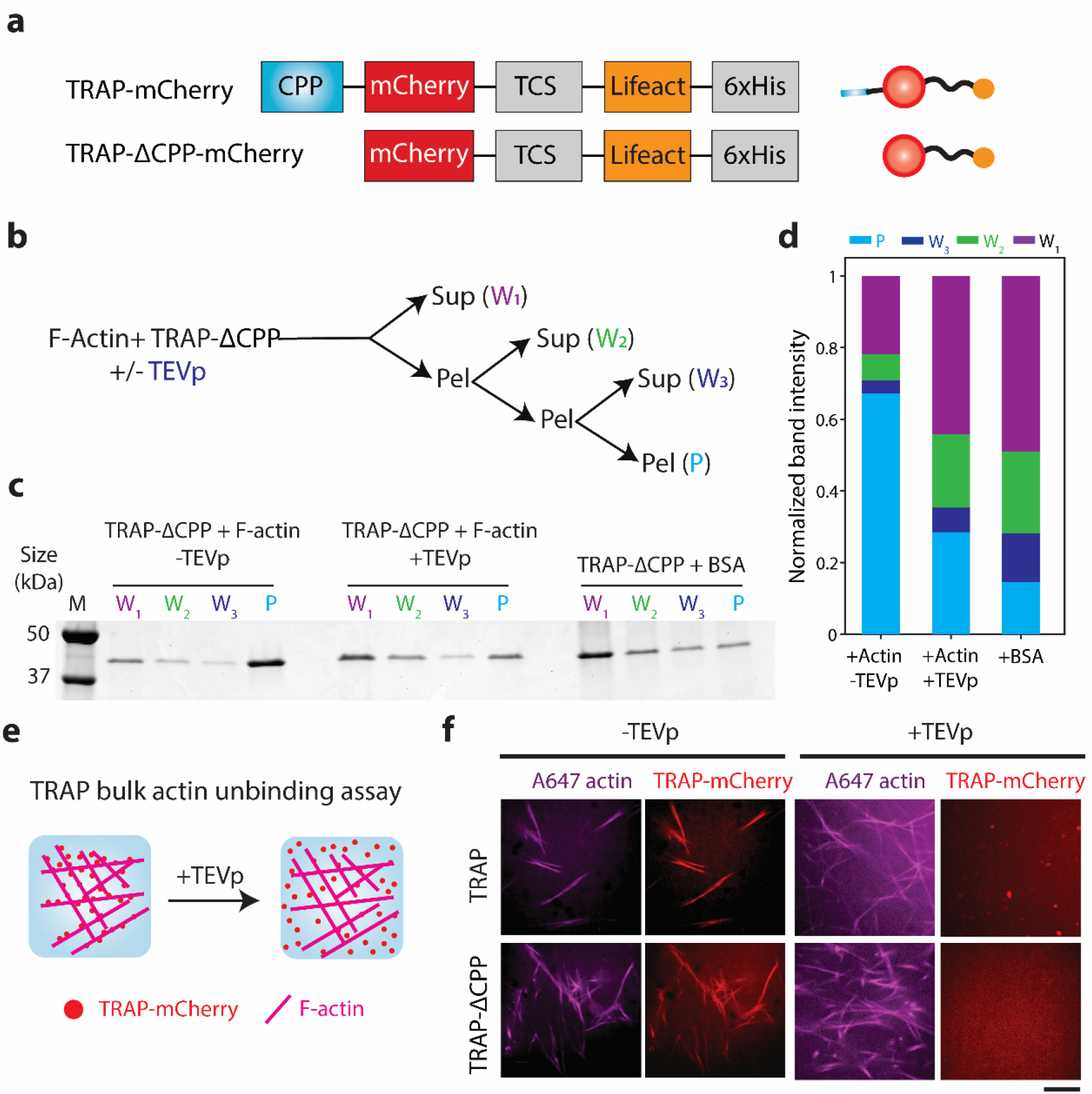
TEVp-mediated unbinding of TRAP-mCherry from actin bundle. **a**. Schematics of constructs of TRAP-mCherry and TRAP-ΔCPP-mCherry (left) and their corresponding protein depictions (right). **b**. Flow chart of the F-actin and TRAP-ΔCPP-mCherry co-pelleting assay. **c**. Representative SDS-PAGE gel image of the pellet and supernatant fractions from the co-pelleting assay of 1 µM TRAP-ΔCPP-mCherry and 5.5 µM of preformed actin incubated for 30 min followed by treatment of 10 units of TEVp. BSA was used as a control instead of F-actin. Lane M indicates the protein ladder. **d**. Bar graph of the normalized band intensities quantified from the SDS-PAGE presented in panel **c**. For each protein, band intensities were normalized by the sum of intensities in all lanes for the corresponding protein. **e**. Illustrations of fluorescence microscopy-based unbinding assay of TRAP-mCherry from F-actin in the presence of TEVp. **f**. Representative confocal fluorescence microscopy images of TRAP-mCherry or TRAP-ΔCPP-mCherry (red) and F-actin (magenta) with or without TEVp treatment for 30 min. Scale bar: 10 μm.

To assess the actin-binding capability of TRAP-mCherry and TRAP-ΔCPP-mCherry, we performed two different assays. First, we conducted a co-pelleting assay of filamentous actin and TRAP-ΔCPP-mCherry. In the absence of membrane in this assay, we reasoned that the actin-binding capability of TRAP-mCherry would be similar to its control variant lacking the CPP domain. The co-pelleting assay of an initial volume of 40 µL consisted of three consecutive ultracentrifugation steps followed by supernatant recovery (20 µL) and pellet (20 µL) resuspension in their original volumes (*i*.*e*., 40 µL) before the next ultracentrifugation run (**Figure 2b**). The entangled filamentous actin is expected to enrich in the pellet fraction while small soluble molecules remain distributed in both pellet and supernatant fractions equally. Therefore, if TRAP tightly binds to actin, it would be enriched in the pellet fraction (P) compared to the supernatant wash fractions (W1, W2, W3).

Consistent with this, we found TRAP-ΔCPP-mCherry was enriched in the pellet fraction only when it contained its actin-binding domain in the absence of TEVp (**Figure 2c, d**), as assessed by SDS-PAGE analysis. In the presence of TEVp or when incubated with a non-binding protein like BSA, TRAP did not enrich in the pellet fraction, and the co-pelleting assay behaved as serial 2-fold dilutions.

In addition to the co-pelleting assay, we imaged TRAP-actin binding and TEVp-mediated TRAP detachment from F-actin using confocal fluorescence microscopy. We reasoned that mixing fluorescently labeled F-actin with TRAP-mCherry would reveal TRAP-mCherry binding to F-actin through colocalization of F-actin and TRAP-mCherry (**Figure 2e**). The addition of TEVp to the solution will then detach the TRAP-mCherry from actin and preclude the colocalization of F-actin and mCherry signals. Indeed, TRAP-mCherry and TRAP-ΔCPP-mCherry colocalized with F-actin in the absence of TEVp (**Figure 2f**). In the presence of TEVp, however, both TRAP-mCherry and TRAP-ΔCPP-mCherry were uniformly dispersed across the field of view, suggesting their detachment from F-actin due to the loss of their Lifeact domain mediated by TEVp cleavage. These results demonstrate TEVp-mediated detachment of TRAP-mCherry from F-actin, thereby validating the use of F-actin as a soluble, large scaffold for sequestering the cargo.

### 2.3 TRAP-mCherry release from GUVs is mediated by TEVp and CPP

Following the confirmation of TRAP-mCherry sequestration by actin, we next hypothesized that the tight binding between TRAP and actin would retain the TRAP when encapsulated in GUVs, thus inhibiting it from freely diffusing out. Additionally, given the complete obliteration of colocalization of actin and TRAP signals in the presence of TEVp, we reasoned that the presence of TEVp in the GUV would drive TRAP unbinding from actin and induce its secretion (**Figure 3a**).

**Figure 3.**
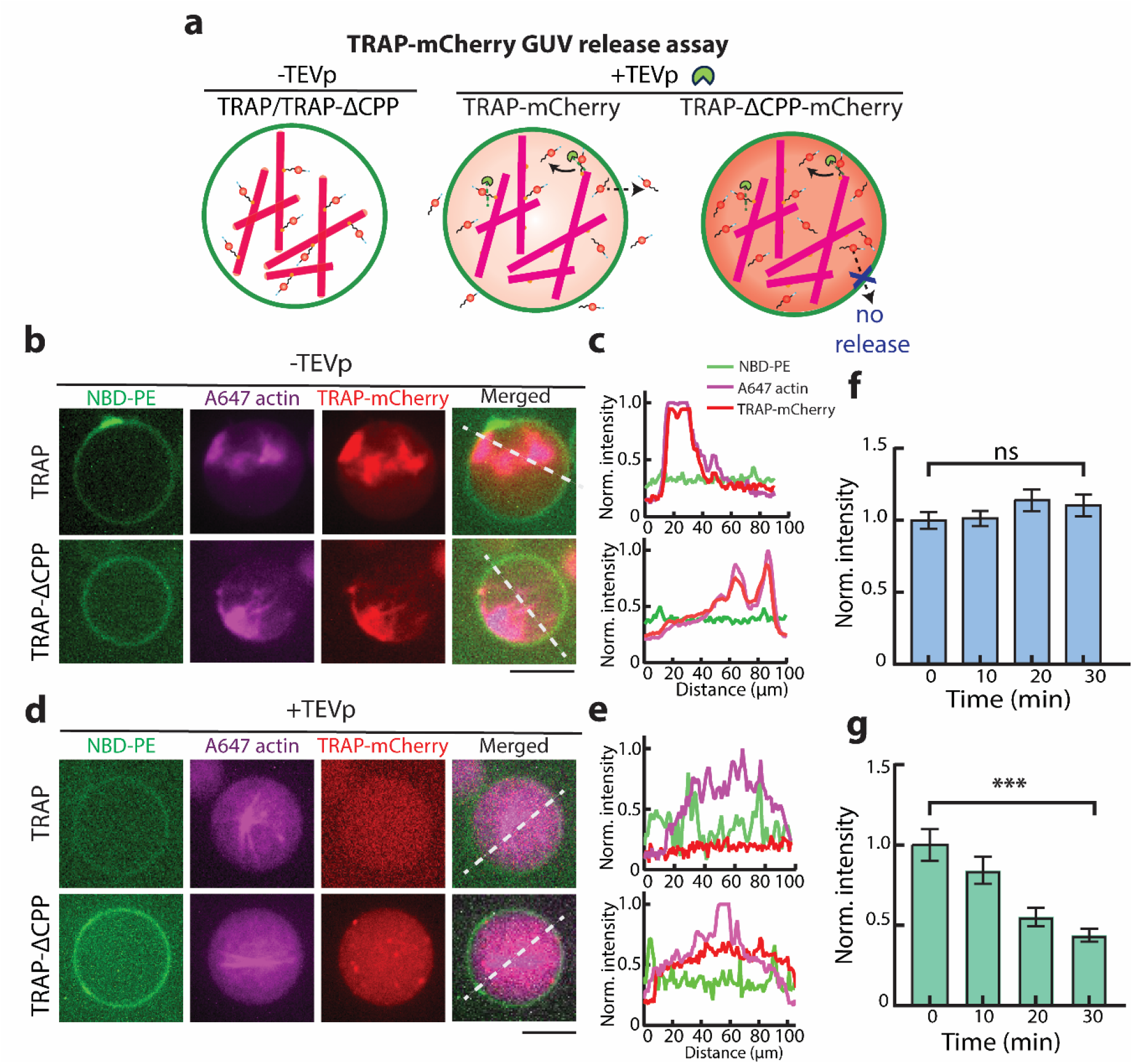
TEVp-mediated TRAP-mCherry release from GUVs. **a**. Schematic illustrations show the binding of TRAP-mCherry and TRAP-ΔCPP-mCherry to F-actin in GUVs in the absence of TEVp. Introduction of TEVp dissociates TRAP and TRAP-ΔCPP from F-actin, leading to the release of TRAP-mCherry, but not TRAP-ΔCPP-mCherry. **b**. Confocal fluorescence microscopy images of GUVs (green) encapsulating TRAP-mCherry or TRAP-ΔCPP-mCherry (red) and F-actin (magenta) in the absence of TEVp. **c**. Normalized fluorescence intensity profiles of NBD-PE (green), actin (red), TRAP-mCherry (magenta) along the white line in the merged channel in panel **b**. The legend shows the fluorescence signal each profile color represents in panels **c** and **e. d**. Representative confocal fluorescence microscopy images of GUVs (green) showing the unbinding of TRAP-mCherry/TRAP-ΔCPP-mCherry (red) from F-actin (magenta) in the presence of TEVp. **e**. Normalized fluorescence intensity profiles of NBD-PE (green), actin (red), TRAP-mCherry (magenta) along the white line in the merged channel in panel **e. f**. Time course fluorescence intensity of TRAP-ΔCPP-mCherry in GUVs in the presence of TEVp. ns = non-significant. Error bars represent the standard error of the means. (*n* = 50 vesicles, three experiments). **g**. Time course fluorescence intensity change of TRAP-mCherry in GUVs in the presence of TEVp. *** represents *p* < 0.001. Error bars represent the standard error of the means. (*n* = 50 vesicles, three experiments). Scale bar: 10 μm.

To test this hypothesis, we encapsulated TRAP-mCherry along with F-actin in GUVs made from DOPC and cholesterol. Consistent with our *in vitro* bulk experiments, we observed strong colocalization of both TRAP-mCherry and TRAP-ΔCPP-mCherry with the actin network within GUVs (**Figure 3b**). Fluorescence intensity profile analysis confirmed the colocalization of mCherry and labeled actin signals, indicating the tight TRAP-actin binding is sustained when encapsulated in GUVs (**Figure 3c**).

When TEVp was co-encapsulated along with TRAP-mCherry and actin, we did not observe colocalization of mCherry and actin signals (**Figure 3d**), which was confirmed by intensity profile analysis (**Figure 3e**). Additionally, while the average intensity of TRAP-ΔCPP-mCherry within GUVs remained constant over time (**Figure 3f**), the fluorescence intensity of TRAP-mCherry gradually decreased over 30 min (**Figure 3g**). This decrease suggests that the detachment of TRAP-mCherry from actin by TEVp enabled CPP-mediated translocation of TRAP-mCherry across the bilayer, effectively allowing it to be secreted from GUVs. The constant fluorescence intensity of TRAP-ΔCPP-mCherry underscores that although it was unbound from actin, the lack of the CPP domain inhibited its secretion. Collectively, these results demonstrate that the inclusion of CPP and Lifeact in the TRAP construct allows TEVp-mediate cargo (*e*.*g*., mCherry) secretion from GUVs.

### 2.4 Light-activated TRAP release demonstrates TRAP modularity

Given the successful TEVp-mediated TRAP secretion from GUVs, we reasoned that linking TEVp activity to some external stimulus could make cargo secretion triggerable. UV light-mediated release of fluorescein di-β-D-galactopyranoside and TEVp from small unilamellar vesicles (SUVs) doped with 1,2-bis(10,12-tricosadiynoyl)-sn-glycero-3-phosphocholine (DC(8,9)PC) were recently demonstrated by Hindley *et al*.^[53]^ and Sihorwala *et al*.,^[34]^ respectively. DC(8,9)PC is a UV-responsive lipid that undergoes photopolymerization upon UV (λ = 254 nm) irradiation, which leads to pore formation in the SUV bilayer. Therefore, we hypothesized that UV-activated protease release from DC(8,9)PC SUVs encapsulating TEVp (denoted as DC(8,9)PC{TEVp}) would induce TRAP secretion by detaching it from the bound actin network (**Figure 4a**).

**Figure 4.**
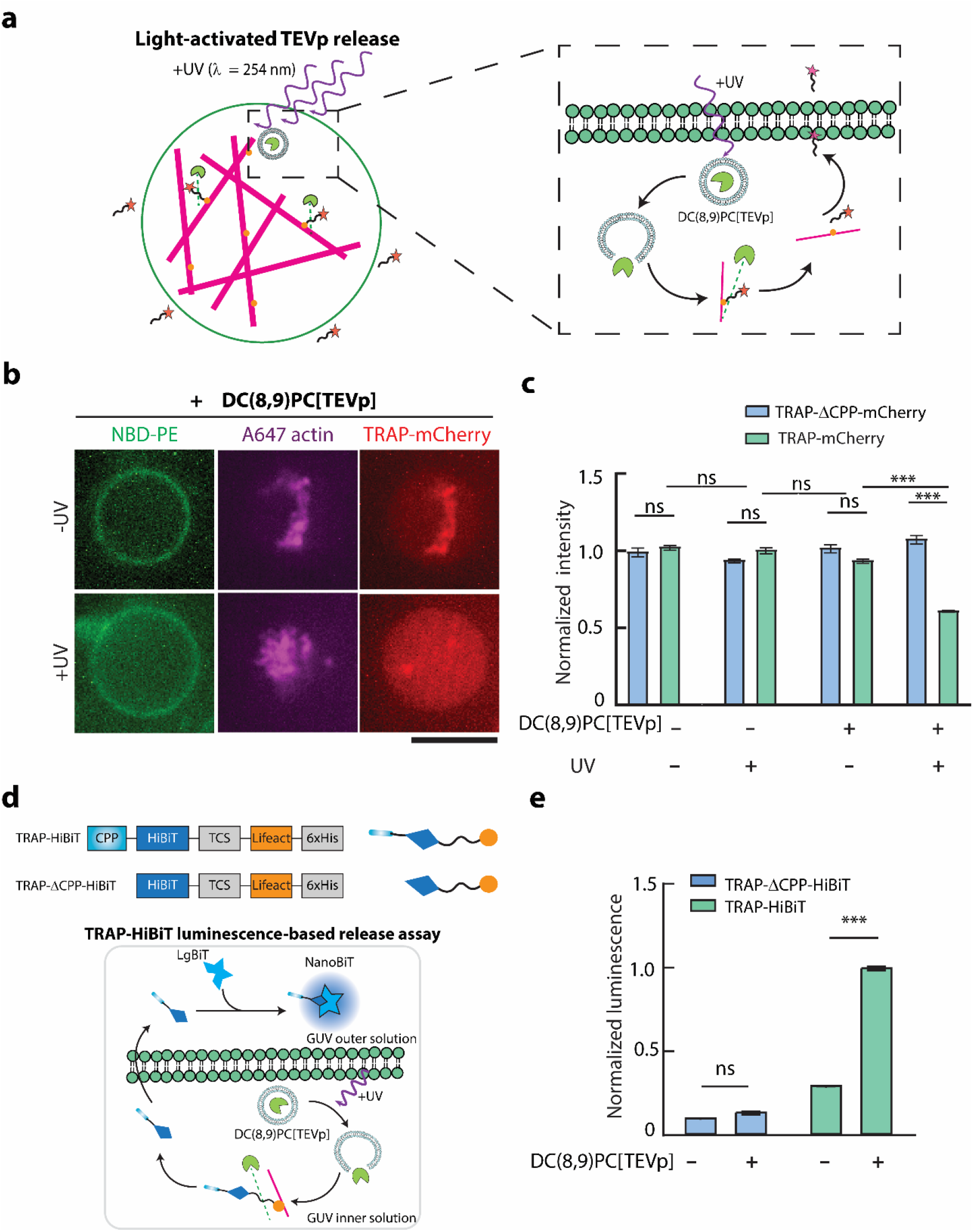
Light-activated TEVp-mediated cargo release from GUVs. **a**. Schematic representation of UV (λ= 254 nm)-activated TEVp release from DC(8,9)PC liposomes (depicted as DC(8,9)PC-{TEVp}) encapsulated in GUVs which induces cargo release from the GUVs through cleavage of TCS on TRAP. Inset shows detailed steps of UV-triggered release of TEVp, cleaving TRAP and allowing the CPP-cargo to traverse the GUV membrane **b**. Representative confocal fluorescence microscopy images of GUVs (green) encapsulating F-actin (magenta) and sTRAP-mCherry (red) with or without 10 min exposure to UV light. Scale bar: 10 μm. **c**. Bar graphs comparing the mCherry fluorescence intensity in GUVs encapsulating TRAP-mCherry in the presence or absence of DC(8,9)PC{TEV} liposomes and UV illumination. Error bars represent standard error of the means, *n* > 50 from 3 independent trials. **d**. Top schematic shows the constructs of TRAP-HiBiT and TRAP-ΔCPP-HiBiT and their corresponding protein depictions. Bottom shows the schematic illustration of TRAP-HiBiT luminescence assay based on Promega Nano-Glo assay. **e**. Normalized luminescence readout from NanoBiT formed by LgBiT dimerization with TRAP-HiBiT or TRAP-ΔCPP-HiBiT released from GUVs in the absence of presence of DC(8,9)PC{TEVp} liposomes. Error bars represent the standard error of the means (*n*= 50 vesicles per condition from 3 independent experiments). ^***^ represents *p* < 0.001. n.s. stands for not significant.

We prepared UV-responsive DC(8,9)PC{TEVp} SUVs and encapsulated them within GUVs along with each TRAP variant as well as F-actin. To test whether DC(8,9)PC{TEVp} SUVs would trigger cargo release through cleavage of the TCS on TRAP, we observed encapsulated TRAP-mCherry release from GUVs exposed to UV light. Confocal fluorescence microscopy of TRAP-mCherry revealed a loss of mCherry localization with actin and a decrease in average mCherry intensity in GUVs exposed to UV light (**Figure 4b**). In contrast, GUVs encapsulating TRAP-mCherry incubated in the dark maintained strong colocalization of mCherry and actin, consistent with a lack of protease activity. Quantification of mCherry fluorescence intensity in GUVs confirmed that TRAP-mCherry was released due to its CPP domain and only in the presence of both DC(8,9)PC{TEVp} SUVs as well as UV irradiation (**Figure 4c**). Consistent with our earlier results, encapsulation of DC(8,9)PC{TEVp} SUVs along with TRAP-ΔCPP-mCherry in GUVs exposed to UV resulted in unbinding of TRAP-ΔCPP-mCherry from actin but not its release (**Figure S2**). These results demonstrate that TRAP secretion from synthetic cells can be triggered by an external physical stimulus that drives TEVp release and activation within GUVs.

To demonstrate the modularity of TRAP platform, we switched its cargo from a fluorescent protein, mCherry, to HiBiT, a short peptide that binds with high affinity to a larger subunit LgBiT to reconstitute a functional luciferase NanoBiT. NanoBiT emits blue light in the presence of its substrate, furimazine, which can be captured with a plate reader. We hypothesized that if HiBiT is encapsulated within GUVs and LgBiT resides in the extracellular environment, they would not be able to form NanoBiT unless HiBiT was secreted from GUVs using the TRAP platform. Thus, we created and purified constructs TRAP-HiBiT and TRAP-ΔCPP-HiBiT by replacing mCherry with HiBiT (**Figure 4d**). To make purification of TRAP-HiBiT and TRAP-ΔCPP-HiBiT proteins through affinity purification facile, we included a soluble tag between the TEVp cleavage site and the Lifeact domain. Specifically, we used a truncated mCherry lacking the 11^th^ β-sheet as a ‘dark mCherry’ soluble tag.^[54]^ This allowed TRAP-HiBiT to have a similar size to TRAP-mCherry, thus simplifying its purification.

Our bulk luminescence measurements confirmed the lack of luciferase activity from both LgBiT and HiBiT on their own as well as high luminescence signals when TRAP-HiBiT and TRAP-ΔCPP-HiBiT with LgBiT were mixed (**Figure S3**). To test TRAP-HiBiT secretion, we encapsulated TRAP-HiBiT along with F-actin and DC(8,9)PC{TEVp} SUVs. After UV irradiation, we added LgBiT to the outer solution of GUVs and incubated the solution to allow NanoBiT formation. Upon addition of furimazine, we measured a strong luminescence signal, suggesting successful HiBiT secretion and thereby dimerization with LgBiT (**Figure 4e**). When we repeated this experiment with TRAP-ΔCPP-HiBiT or without DC(8,9)PC{TEVp} SUVs, we did not detect a strong luminescence signal. Therefore, the inclusion of DC(8,9)PC{TEVp} and the CPP domain is essential, collectively indicating the role of TEVp in initiating CPP-mediated TRAP-HiBiT secretion to the extracellular environment. Taken together, these results demonstrate the modularity of TRAP, which enabled the secretion of two different cargos as well as its ability to accommodate stimuli-driven secretion from synthetic cells.

## 3 Discussion

We designed and demonstrated a novel protein-based platform, termed TRAP, which facilitates triggerable, TEVp-based, and CPP-mediated selective protein secretion from synthetic cells. Since CPP allows passive diffusion of cargo outside the GUV, TRAP was designed to be sequestered by an intracellular actin network. This design ensures that secretion occurs only when TRAP is detached from the actin network via TEVp cleavage. Utilizing SUVs containing the UV-sensitive lipid DC(8,9)PC, we successfully demonstrated UV-triggered TEVp release inside GUVs, resulting in TRAP secretion. To highlight the modularity of the TRAP platform, we demonstrated UV-driven secretion of two distinct cargos by merely swapping the cargo domain of TRAP.

A key aspect of the TRAP platform is its ability to function as a purified protein, bypassing the need for synthesis via a cell-free expression system, as others have previously shown.^[45,46]^ This capability enabled rapid secretion, observable within 30 min. Additionally, with TRAP sequestered to the intracellular actin network in the absence of protease activity, its secretion can be quickly and efficiently triggered by rapid protease activation.

As TRAP release is dependent on the proteolytic cleavage of the sequence connecting the cargo to the actin-binding domain, the TRAP platform can accommodate a variety of inputs provided they enable TEVp activation. For example, blue-light mediated split TEVp reconstitution,^[55]^ rapamycin-induced TEVp activation,^[56]^ or inducible TEVp expression using a cell-free expression system can be designed to trigger secretion. The flexibility of TRAP in accommodating different inputs and its modular design in allowing various cargos make it a powerful tool for biosensing applications.

By incorporating different protease cleavage sites between the cargo and actin-binding domains, it is possible to engineer multiple TRAP variants, each used for the secretion of a specific cargo, thus enabling independent secretion of multiple cargos. Proteases that precisely cleave their corresponding unique cleavage sites such as TEV, TVMV, thrombin, and HRV3C are ideal candidates for multiplexed TRAP-mediated cargo secretion. Furthermore, tandem placement of different protease cleavage sites instead of a single cleavage sequence allows for encoding an OR gate for TRAP secretion, where any input that drives separation of cargo molecules from the actin-binding domain would lead to secretion. This modularity of TRAP, enabling multiple cargos while allowing the release of each cargo contingent on receiving a unique input, makes it a versatile tool for programming combinatorial sensing in synthetic cells for therapeutic and engineering applications.

In addition to sensing applications, TRAP promises many possibilities in engineering intercellular synthetic cell and synthetic cell-natural cell communication. For example, we demonstrated a split luciferase activation in the extracellular environment through TRAP-HiBiT release. Inclusion of LgBiT in a second population of synthetic cells could enable TRAP-mediated intercellular communication between secretory sender synthetic cells that release TRAP-HiBiT and receiver synthetic cells that encapsulate LgBiT and become actively bioluminescent upon receiving TRAP-HiBiT. Furthermore, the release of an orthogonal protease from one population of synthetic cells can prompt TRAP-mediated release of another orthogonal protease from a second population of synthetic cells, thereby allowing the encoding of cascaded TRAP-based secretory reactions across different synthetic cell populations. Lastly, when synthetic cells are mixed with natural cells, TRAP may facilitate communication between synthetic cells and natural cells. For instance, the secretion of DNA-editing enzymes like Cre recombinase or a Cas protein can induce transcriptional regulation in living receiver cells. Therefore, TRAP could find use in engineering smart biomaterials and therapeutic protein delivery applications.

In summary, this work marks a significant advancement in the design of biomimetic synthetic systems equipped with controlled, multiplexed secretion capabilities. As the field of synthetic cell engineering evolves, future developments could focus on stimuli-responsive biomaterials^[57]^ as well as biosensing systems with TRAP as their central output-generating secretory mechanism. Furthermore, integrating synthetic cells equipped with different variants of TRAP with living cells could enable triggerable targeted cargo release, allowing synthetic cells to interact with and modulate living cells for therapeutic purposes.

## 4 Experimental Section

### 4.1 Cloning and DNA preparation

DNA sequences encoding for TRAP-mCherry, TRAP-ΔCPP-mCherry, TRAP-HiBiT, and TRAP-ΔCPP-HiBiT flanked by a Shine-Dalgarno ribosome binding sequence and a 6xHis tag on their 5’ and 3’ ends, respectively, were synthesized and cloned into a pET24(+) vector by Twist Bioscience. The details of the DNA sequences for each construct are provided in Supplementary Information.

### 4.2 Proteins and reagent

Both TRAP-mCherry and TRAP-HiBiT proteins along with their control variants lacking the CPP domain were purified using conventional His-tagged protein purification methods described elsewhere.^[54,58]^ Single colonies of BL21(DE3)pLysS cells transformed with the plasmid encoding the protein of interest were picked from LB agar plates and were grown in 5 mL LB media supplemented with 50 μg/mL kanamycin overnight in a shaker incubator at 37^°^C shaking at 220 rpm. Next, the culture was diluted in 1 L LB containing 50 μg/mL kanamycin and the cells were grown at 37^°^C shaking at 220 rpm until the OD_600_ reached 0.6. The cells were then induced with 0.1 mM isopropyl β-d-1-thiogalactopyranoside (IPTG) and incubated overnight at room temperature shaking at 220 rpm. Cells were then harvested by centrifugation at 5000 g for 15 min, and the pellet was resuspended in 30 mL lysis buffer containing 20 mM Tris pH 8.0, 500 mM NaCl, 5 mM imidazole, 1 mM 4-(2-aminoethyl)benzenesulfonyl fluoride hydrochloride (AEBSF, Sigma-Aldrich), and two tablets of protease inhibitor cocktails (Roche). Cells were then lysed using a tip sonicator (Branson Sonifier 450) and the lysate was centrifuged at 30,000 g for 30 min at 4°C. Next, the supernatant was loaded onto a 1 mL HisTrap FF column (Cytiva) mounted on an AKTA Start fast protein liquid chromatography (FPLC) system. The column was then washed with 15 mL of washing buffer containing 20 mM Tris pH 8.0, 500 mM NaCl, and 25 mM imidazole. Lastly, individual 1 mL fractions were eluted by passing elution buffer (20 mM Tris pH 8.0, 500 mM NaCl, and 250 mM imidazole) through the column. The purification quality was assessed by SDS-PAGE and fractions with the highest concentration of purified protein were pooled and dialyzed against dialysis buffer (20 mM Tris pH 8.0, 25 mM NaCl, and 1 mM EDTA) overnight at 4^°^C. Next, protein concentration was measured using NanoDrop (extinction coefficient predicted by Benchling biochemical analysis tool) and individual aliquots were made and stored at -80^°^C until use.

Unlabeled actin and fluorescent labeled ATTO 488 actin were purchased from Cytoskeleton Inc, USA. Fascin was purified from *E. coli* as a glutathione-S-transferase (GST) fusion protein following the protocols described elsewhere.^[12,59]^ For purification, BL21(DE3) cells were transformed with pGEX-6P-1 fascin plasmid encoding the sequences of GST translationally fused to fascin by a PreScission protease cleavage site. Cells were grown at 37°C while shaking at 220 rpm, until the OD_600_ reached 0.5–0.6. Protein expression was induced with 0.1 mM IPTG, and the cell culture was incubated at 24°C for 8 h. Cells were harvested by centrifugation at 4,000 *g* for 15 min and washed with PBS once. The cell pellet was resuspended in lysis buffer (20 mM K-HEPES, pH 7.5, 100 mM NaCl, 1 mM EDTA, and 1 mM PMSF) and ruptured by sonication. The lysate was then centrifuged at 45,000 *g* for 25 min and the supernatant was loaded on a 1 mL GSTrap FF column (Cytiva) using an AKTA Start FPLC system. The column was washed with 15 mL washing buffer (20 mM K-HEPES, pH 7.5, 500 mM NaCl). Next, 2 mL of cleavage buffer (50 mM Tris-HCl, pH7.5, 150 mM NaCl, 1 mM EDTA, and 1 mM dithiothreitol) containing 50 units of PreScission protease (GenScript) was loaded on the column and incubated overnight at 4°C to cleave GST from fascin. The GST-free fascin was then eluted with 5 mL of elution buffer (PBS). Purified products were dialyzed against 1 L of PBS twice for 3 h and once overnight at 4°C. Protein concentration was measured by NanoDrop. The protein concentration was adjusted using a Centricon filter (Millipore), and the concentrated protein was aliquoted and stored at -80.

### 4.3 *In vitro* actin-binding assay

5.5 μM actin was polymerized by incubating in polymerization buffer (50 mM KCl, 2 mM MgCl_2_, 0.2 mM CaCl_2_, and 4.2 mM ATP in 15 mM Tris, pH 8.0) for 15 min at room temperature. 2.5 μM of fascin was incubated with preformed actin filaments for 10 min at room temperature to prepare the fascin-bundled actin. A flow cell was prepared using two glass cover slides of 24 mm by 50 mm and 18 mm by 18 mm adhered together by two double-sided tapes. 10 μL of 0.1 mg/ml BSA solution in G-buffer (5 mM Tris, pH 8.0 and 0.2 mM CaCl_2_) was loaded to the flow cell. After incubating for 5 min, 10 μL fascin-bundled actin was introduced to the flow channel. After incubating for 5 min, flow channel was washed with G-buffer twice. Then 10 μL of 1 μM TRAP-mCherry was added to the flow channel. After 10 min incubation, flow channel was washed with 10 μL of G-buffer to remove the unbound TRAP-mCherry before imaging. For the experiments with TEVp, TRAP-mCherry was incubated with 10 units of TEVp for 30 min before addition to the flow channel prior to imaging.

### 4.4 Actin-TRAP-mCherry co-pelleting assay and SDS-PAGE analysis

Preformed actin was incubated with 1μM of TRAP-ΔCPP-mCherry in polymerization buffer for 30 min with or without TEVp. BSA was used as a control instead of actin and incubated with TRAP-ΔCPP-mCherry for 30 min. Then 40 μL of each sample was transferred to an ultracentrifuge tube (Beckman Coulter) and centrifuged at around 100,000 g for 15 min at room temperature using an Airfuge (Beckman Coulter). After centrifugation, 20 μL of the supernatant (W1) was carefully recovered so as not to disturb the pellet and was transferred to a 1.5 mL microcentrifuge tube. The remaining 20 μL was mixed with 20 μL polymerization buffer and the pellet was resuspended by gently pipetting up and down. This process was repeated two more times to obtain supernatant fractions W2 and W3. The final remaining 20 μL of pellet was resuspended and stored as the pellet fraction (P). W1, W2, W3, and P were mixed with 4x Laemmli loading buffer (Bio-Rad) containing 10% 2-mercaptoethanol. Samples were heated at 90°C for 10 min before loading on a 4−20% Bis-Tris PAGE gel (GeneScript). The gel was stained using SimplyBlue stain (Invitrogen) and imaged by a Sapphire Biomolecular Imager (Azure biosystems) with 658/710 nm excitation/emission wavelengths. The band intensities in each lane for every protein illustrated in **Figure 2c** were measured using ImageJ gel analyzer tool, and the intensity values were normalized by the sum of intensities for each protein.

### 4.5 GUV preparation

To prepare GUVs, 0.4 mM mixture of lipids containing 69.9% 1,2-dioleoyl-sn-glycero-3-phosphocholine (DOPC), 30% cholesterol, and 0.1% 1,2-dioleoyl-sn-glycero-3-phosphoethanolamine-N-(7-nitro-2-1,3-benzoxadiazol-4-yl)_(NBD-PE) in a 4:1 mixture of silicone oil (Sigma-Aldrich) and mineral oil (Sigma-Aldrich) was first made in a glass tube. The lipid/oil mixture was prepared a day before the experiment and stored at 4°C till use. All lipids were purchased from Avanti Polar Lipids.

Next, a reaction mixture of 5.5 μM actin (including 10% ATTO 647 actin) in polymerization buffer was prepared and kept on ice for 15 min. 2.5 μM of fascin and 1 μM of TRAP-mCherry variants (with or without CPP) were then added to the sample followed by addition of 7.5% OptiPrep density gradient medium (Sigma-Aldrich). GUVs were then immediately generated following the cDICE method. A cDICE chamber was 3D-printed with Clear V4 resin (Formlabs) using a Form 3 3D printer (Formlabs) and mounted on the rotor of a benchtop stirrer hot plate and rotated at 1,200 rpm. Around 0.7 mL of aqueous outer solution (230 mM D-glucose matching the osmolarity of the inner solution) and around 5 mL of lipid-in-oil dispersion were sequentially transferred into the rotating chamber. To generate GUVs, a water-in-oil emulsion was first created by vigorously pipetting 20 μL of reaction mix in 0.7 mL of lipid-in-oil mixture and the emulsion was pipetted into the rotating chamber. The chamber was unmounted after 30 s of rotation following the addition of water-in-oil emulsion droplets, and the GUVs were collected by recovering the aqueous outer solution. The GUVs were transferred to a 96-well plate for imaging.

### 4.6 Preparation of DC(8,9)PC{TEVp} liposomes for light-based activation of TRAP

Light-sensitive liposomes encapsulating TEVp were prepared following the protocol described by Sihorwala *et al*.^[34]^ A 10 mg/mL lipid solution containing 79.5% DPPC, 20% DC(8,9)PC, and 0.5% DSPE-PEG_2000_ lipids in chloroform was prepared and exposed to a gentle stream of argon gas to evaporate the chloroform. The lipids were prepared in the absence of light and covered with aluminum foil. The film was incubated in a desiccator for 2 hours to ensure complete evaporation of residual solvent. Next, the lipid film was hydrated with an aqueous solution containing 100 units of TEVp, 1 mM DTT, 25 mM HEPES, 150 mM NaCl, pH 7.4, incubated at 42°C and vortexed for 30 s. The solution was then extruded 23 times through a 100 nm polycarbonate filter mounted on the Avanti mini-extruder preincubated at 42°C to prepare liposomes encapsulating TEVp. The resulting liposomes were kept at 4°C for 30 minutes. Finally, liposome samples were centrifuged at 16,900 g for 10 min at room temperature to separate any large lipid aggregates from liposomes.

### 4.7 Light-triggered TEVp release in GUVs

20 μL of GUV inner solution was made by addition of 4.7 μL DC(8,9)PC{TEVp} liposomes, 7.5% OptiPrep, and 1 µM TRAP-mCherry or TRAP-HiBiT to a mixture of 5.5 μM actin in polymerization buffer (for TRAP-mCherry, 2.5 μM fascin and 0.6 μM ATTO 647-labeled actin was mixed with actin solution as well) preincubated on ice for 15 min. GUVs were prepared using cDICE described previously. The GUVs containing DC(8,9)PC{TEVp} liposomes and the control samples lacking the liposomes were irradiated with 254 nm UV light for 10 minutes, followed by incubation at 30°C for 1 hour. Simultaneously, aliquots of non-UV irradiated vesicles, prepared with and without liposomes, were also incubated at 30°C for 1 hour. The GUVs encapsulating TRAP-mCherry were then observed using confocal microscopy.

### 4.8 Bioluminescence measurement

Following the production of GUVs containing light-sensitive liposomes and their corresponding control variants, GUVs encapsulating TRAP-HiBiT were pelleted by centrifugation at 300 g for 15 min. The supernatant was removed, and the GUV pellet was resuspended in 50 μL of aqueous outer solution. Then, from each experimental condition (+/- UV and +/- liposomes), 30 μL of GUV solution was transferred into a 96-well half-area well-plate (Corning), and TEVp release was induced by UV irradiation as described in the previous section. Next, 1.5 μL of 20 μM LgBiT was added to each well and the well plate was incubated for 15 min at room temperature to allow NanoBiT formation. Then, 2 μL of furimazine stock (Nano-Glo® assay, Promega) was added to each well, and the plate was immediately shaken manually before the bioluminescence signals were captured every minute for 60 min using a Synergy H1 (BioTek) multimode plate-reader. All measurements were done using 1s integration time and a gain of 130. The luminescence values of the UV and non-UV irradiated samples were compared following the subtraction of luminescence from their respective control conditions.

### 4.9 Confocal fluorescence microscopy imaging

Fluorescence images were captured using an oil immersion 60x/1.4 NA Plan-Apochromat objective on an Olympus IX-81 inverted microscope equipped with a spinning-disk confocal (Yokogawa CSU-X1), solid-state lasers (Solamere Technology) controlled by a National Instrument DAQmx, and an iXON3 EMCCD camera (Andor Technology). Image acquisition was controlled by MetaMorph software (Molecular Devices). Lipid, TRAP-mCherry, and actin fluorescence images were taken with 488-nm laser excitation at an exposure time of 1000 ms, 561-nm laser excitation at an exposure time of 400 ms, and 647-nm laser excitation at an exposure time of 400 ms, respectively. A Semrock quad-band bandpass filter was used as the emission filter.

### 4.10 Image and statistical analyses

All GUV images were analyzed by ImageJ. The intensity profiles presented in **Figures 3c** and **3e** were obtained by measuring the fluorescence intensity along the corresponding indicated lines in **Figures 3b** and **3d**, respectively. Background signal was measured and subtracted from the intensity values followed by signal intensity normalization to the maximum signal. For analyzing fluorescence intensity presented in **Figures 3f** and **3g**, average total fluorescence intensity of TRAP-mCherry with or without CPP for individual GUVs was measured at different time points and normalized to the average signal at *t* = 0.

All the experiments were performed with at least three independent replicates. Statistical analyses and graph generation were performed using GraphPad Prism. The unpaired *t*-test or one-way and two-way ANOVA tests corrected with Tukey’s Honest Significant Difference were used. Calculated *p*-values are presented in **Table S1** in Supporting Information.

## Acknowledgements

This research was funded by the National Institutes of Health R21 AI173559 to A.P.L. and G.D.L., R01 EB030031 to A.P.L., and by the National Science Foundation EF1935265 to A.P.L. The authors received funding from National Institutes of Health R01 CA238402, R33CA225549, U24CA237683, and R37CA222563; National Science Foundation 214104; DOD Breast Cancer Research Program W81XWH2210120; and the W.M. Keck Foundation (G.D.L.). H.M. acknowledges the support from the Rackham Pre-doctoral Fellowship. We thank Mrittika Sarkar for help with image analysis for Figure 4c and the Liu lab for helpful discussion.

## Author Contributions

Conceptualization: H. M., M.A. and A.P.L. Methodology: M. A. and H. M. Investigation: M. A. and H. M. Visualization: M. A. and H. M. Supervision: A.P.L. and G. D. L. Writing—original draft: M. A. and H. M. Writing—review & editing: M. A., H. M., G. D. L. and A.P.L.

## References

[1] C. Guindani, L. C. da Silva, S. Cao, T. Ivanov, K. Landfester, Angewandte Chemie 2022, 134, e202110855.

[2] B. Sharma, H. Moghimianavval, S.-W. Hwang, A. P. Liu, Membranes 2021, 11, 912.

[3] A. Groaz, H. Moghimianavval, F. Tavella, T. W. Giessen, A. G. Vecchiarelli, Q. Yang, A. P. Liu, WIREs Nanomedicine and Nanobiotechnology 2021, 13, e1685.

[4] M. A. Boyd, N. P. Kamat, Trends in Biotechnology 2021, 39, 927.

[5] M. A. Boyd, W. Thavarajah, J. B. Lucks, N. P. Kamat, Science Advances 2023, 9, eadd6605.

[6] A. Zambrano, G. Fracasso, M. Gao, M. Ugrinic, D. Wang, D. Appelhans, A. deMello, T.-Y. D. Tang, Nat Commun 2022, 13, 3885.

[7] B.C. Buddingh’, J. Elzinga, J. C. M. van Hest, Nat Commun 2020, 11, 1652.

[8] H. Moghimianavval, K. J. Loi, S.-W. Hwang, Y. Bashirzadeh, A. P. Liu, 2024.

[9] A. Samanta, L. Baranda Pellejero, M. Masukawa, A. Walther, Nat Rev Chem 2024, 8, 454.

[10] E. Bailoni, M. Partipilo, J. Coenradij, D. A. J. Grundel, D. J. Slotboom, B. Poolman, ACS Synth. Biol. 2023, 12, 922.

[11] S. Berhanu, T. Ueda, Y. Kuruma, Nat Commun 2019, 10, 1325.

[12] Y. Bashirzadeh, S. A. Redford, C. Lorpaiboon, A. Groaz, H. Moghimianavval, T. Litschel, P. Schwille, G. M. Hocky, A. R. Dinner, A. P. Liu, Commun Biol 2021, 4, 1.

[13] J. Powers, Y. Jang, Biomacromolecules 2023, 24, 5539.

[14] B. P. Jena, Current Opinion in Structural Biology 2007, 17, 437.

[15] M. Leabu, G. L. Nicolson, Discoveries 2014, 2, e30.

[16] S. Trikha, E. C. Lee, A. M. Jeremic, The Scientific World Journal 2010, 10, 152590.

[17] T. L. Burgess, R. B. Kelly, Annual Review of Cell and Developmental Biology 1987, 3, 243.

[18] N. I. Md Fadilah, M. S. Mohd Abdul Kader Jailani, M. A. I. Badrul Hisham, N. Sunthar Raj, S. A. Shamsuddin, M. H. Ng, M. B. Fauzi, M. Maarof, J Tissue Eng 2022, 13, 20417314221114273.

[19] K. Francis, B. O. Palsson, Proc Natl Acad Sci U S A 1997, 94, 12258.

[20] E. T. Kavalali, Nat Rev Neurosci 2015, 16, 5.

[21] J. G. Cyster, C. D. C. Allen, Cell 2019, 177, 524.

[22] P. Rorsman, F. M. Ashcroft, Physiological Reviews 2018, 98, 117.

[23] F. M. Gribble, F. Reimann, Nat Rev Endocrinol 2019, 15, 226.

[24] T. Kjeldsen, A. S. Andersen, F. Hubálek, E. Johansson, F. F. Kreiner, G. Schluckebier, P. Kurtzhals, Trends in Biotechnology 2024, 42, 464.

[25] A. E. Vlahos, J. Kang, C. A. Aldrete, R. Zhu, L. S. Chong, M. B. Elowitz, X. J. Gao, Nat Commun 2022, 13, 912.

[26] X. Wang, L. Kang, D. Kong, X. Wu, Y. Zhou, G. Yu, D. Dai, H. Ye, Nat Chem Biol 2024, 20, 432.

[27] K. P. Adamala, D. A. Martin-Alarcon, K. R. Guthrie-Honea, E. S. Boyden, Nature Chem 2017, 9, 431.

[28] L. Lu, J. W. Schertzer, P. R. Chiarot, Lab Chip 2015, 15, 3591.

[29] G. Mohanan, K. S. Nair, K. M. Nampoothiri, H. Bajaj, Chem. Sci. 2020, 11, 4669.

[30] J. M. Thomas, M. S. Friddin, O. Ces, Y. Elani, Chemical Communications 2017, 53, 12282.

[31] T.-Y. D. Tang, D. Cecchi, G. Fracasso, D. Accardi, A. Coutable-Pennarun, S. S. Mansy, A. W. Perriman, J. L. R. Anderson, S. Mann, ACS Synth. Biol. 2018, 7, 339.

[32] C. Monck, Y. Elani, F. Ceroni, Nat Chem Biol 2024, 1.

[33] C. E. Hilburger, M. L. Jacobs, K. R. Lewis, J. A. Peruzzi, N. P. Kamat, ACS Synth. Biol. 2019, 8, 1224.

[34] A. Z. Sihorwala, A. J. Lin, J. C. Stachowiak, B. Belardi, J. Am. Chem. Soc. 2023, 145, 3561.

[35] R. Lentini, S. P. Santero, F. Chizzolini, D. Cecchi, J. Fontana, M. Marchioretto, C. Del Bianco, J. L. Terrell, A. C. Spencer, L. Martini, M. Forlin, M. Assfalg, M. D. Serra, W. E. Bentley, S. S. Mansy, Nat Commun 2014, 5, 4012.

[36] Ö. D. Toparlak, J. Zasso, S. Bridi, M. D. Serra, P. Macchi, L. Conti, M.-L. Baudet, S. S. Mansy, Science Advances 2020, 6, eabb4920.

[37] X. Wang, L. Tian, H. Du, M. Li, W. Mu, B. W. Drinkwater, X. Han, S. Mann, Chem. Sci. 2019, 10, 9446.

[38] S.-W. Hwang, C.-M. Lim, C. T. Huynh, H. Moghimianavval, N. A. Kotov, E. Alsberg, A. P. Liu, Angewandte Chemie International Edition 2023, 62, e202308509.

[39] G. Chen, R. Levin, S. Landau, M. Kaduri, O. Adir, I. Ianovici, N. Krinsky, O. Doppelt-Flikshtain, J. Shklover, J. Shainsky-Roitman, S. Levenberg, A. Schroeder, Proceedings of the National Academy of Sciences 2022, 119, e2207525119.

[40] D. Fasshauer, Biochimica et Biophysica Acta (BBA) - Molecular Cell Research 2003, 1641, 87.

[41] A. M. Liaci, F. Förster, International Journal of Molecular Sciences 2021, 22, 11871.

[42] Z. Chen, J. Wang, W. Sun, E. Archibong, A. R. Kahkoska, X. Zhang, Y. Lu, F. S. Ligler, J. B. Buse, Z. Gu, Nat Chem Biol 2018, 14, 86.

[43] N. L. Mora, A. L. Boyle, B. J. van Kolck, A. Rossen, Š. Pokorná, A. Koukalová, R. Šachl, M. Hof, A. Kros, Sci Rep 2020, 10, 3087.

[44] Y.-Y. Hsu, S. J. Chen, J. Bernal-Chanchavac, B. Sharma, H. Moghimianavval, N. Stephanopoulos, A. P. Liu, Chem. Commun. 2023, 59, 8806.

[45] N. Krinsky, M. Kaduri, A. Zinger, J. Shainsky-Roitman, M. Goldfeder, I. Benhar, D. Hershkovitz, A. Schroeder, Advanced Healthcare Materials 2018, 7, 1701163.

[46] J. M. Heili, K. Stokes, N. J. Gaut, C. Deich, J. Sharon, T. Hoog, J. Gomez-Garcia, B. Cash, M. R. Pawlak, A. E. Engelhart, K. P. Adamala, Cell Systems 2024, 15, 49.

[47] D. Derossi, A. H. Joliot, G. Chassaing, A. Prochiantz, Journal of Biological Chemistry 1994, 269, 10444.

[48] A. Walrant, L. Matheron, S. Cribier, S. Chaignepain, M.-L. Jobin, S. Sagan, I. D. Alves, Analytical Biochemistry 2013, 438, 1.

[49] P. E. G. Thorén, D. Persson, M. Karlsson, B. Nordén, FEBS Letters 2000, 482, 265.

[50] L. N. Patel, J. L. Zaro, W.-C. Shen, Pharm Res 2007, 24, 1977.

[51] S. T. Henriques, M. N. Melo, M. A. R. B. Castanho, Biochemical Journal 2006, 399, 1.

[52] J. Riedl, A. H. Crevenna, K. Kessenbrock, J. H. Yu, D. Neukirchen, M. Bista, F. Bradke, D. Jenne, T. A. Holak, Z. Werb, M. Sixt, R. Wedlich-Soldner, Nat Methods 2008, 5, 605.

[53] J. W. Hindley, Y. Elani, C. M. McGilvery, S. Ali, C. L. Bevan, R. V. Law, O. Ces, Nat Commun 2018, 9, 1093.

[54] H. Moghimianavval, C. Patel, S. Mohapatra, S.-W. Hwang, T. Kayikcioglu, Y. Bashirzadeh, A. P. Liu, T. Ha, Small 2023, 19, 2202104.

[55] M. Cui, S. Lee, S. H. Ban, J. R. Ryu, M. Shen, S. H. Yang, J. Y. Kim, S. K. Choi, J. Han, Y. Kim, K. Han, D. Lee, W. Sun, H.-B. Kwon, D. Lee, Nat Chem Biol 2024, 20, 353.

[56] P. Renna, C. Ripoli, O. Dagliyan, F. Pastore, M. Rinaudo, A. Re, F. Paciello, C. Grassi, Bioengineering & Translational Medicine 2022, 7, e10292.

[57] A. P. Liu, E. A. Appel, P. D. Ashby, B. M. Baker, E. Franco, L. Gu, K. Haynes, N. S. Joshi, A. M. Kloxin, P. H. J. Kouwer, J. Mittal, L. Morsut, V. Noireaux, S. Parekh, R. Schulman, S. K. Y. Tang, M. T. Valentine, S. L. Vega, W. Weber, N. Stephanopoulos, O. Chaudhuri, Nat. Mater. 2022, 21, 390.

[58] H. Moghimianavval, S. Mohapatra, A. P. Liu, in Mammalian Synthetic Systems (Eds: F. Ceroni, K. Polizzi), Springer US, New York, NY, 2024, pp. 43–58.

[59] Y. Bashirzadeh, H. Moghimianavval, A. P. Liu, iScience 2022, 25, DOI 10.1016/j.isci.2022.104236.

