## Supplemental information for "Light-triggered protease-mediated release of actin-bound cargo from synthetic cells"

**TRAP constructs**

**TRAP-mCherry:**

Ribosome-binding site

Cell-penetrating peptide

mCherry

GGGSG linker

TEVp cleavage site

Lifeact

6xHis tag

AGAAAGTAAGGAGGTTTTTTATGCGTCAAATAAAGATTTGGTTTCAAAATCGTCGGATGAAGT
GGAAGAAGGTAAGCAAGGGTGAAGAAGACAATATGGCAATCATAAAAGAGTTCATGCGGTTT
AAAGTTCACATGGAAGGTTCGGTTAATGGACATGAATTTGAGATTGAAGGTGAAGGGGAAG
GTCGGCCATACGAGGGTACCCAAACTGCTAAGCTCAAAGTCACTAAAGGCGGACCTCTTCC
CTTTGCTTGGGATATTCTTTCTCCGCAATTCATGTATGGTAGTAAAGCCTATGTGAAACATCC
CGCGGACATCCCTGATTATTTAAAGCTTTCCTTTCCGGAAGGCTTTAAATGGGAGCGCGTTA
TGAACTTCGAAGACGGTGGGGTTGTGACGGTGACGCAAGATTCATCCCTGCAAGATGGGG
AGTTCATTTACAAAGTAAAGCTCAGAGGGACGAATTTTCCTTCTGATGGTCCGGTTATGCAA
AAGAAGACGATGGGATGGGAAGCATCGTCCGAGAGAATGTACCCGGAAGATGGGGCTCTT
AAAGGCGAGATTAAACAACGCCTTAAACTTAAAGACGGCGGCCACTACGACGCGGAAGTGA
AGACTACGTACAAAGCAAAGAAGCCAGTACAATTACCGGGAGCCTATAATGTTAACATAAAGT
TGGACATCACTAGTCACAATGAGGATTACACGATTGTTGAGCAATATGAACGCGCAGAAGGA
CGCCACTCAACTGGTGGAATGGACGAGCTGTATAAAGGTGGCGGCGGTTCCGGAGAGAAT
TTATATTTCCAAGGTGGCGGGGGGGGATCGGGTGGTGTTGCGGATTTAATAAAGAAATTTGA
ATCTATTAGTAAAGAAGAAGGCGGCGGCGGTTCAGGGCATCATCATCATCATCATTAA

**TRAP-ΔCPP-mCherry:**

Ribosome-binding site

mCherry

GGGSG linker

TEVp cleavage site

Lifeact

6xHis tag

AGAAAGTAAGGAGGTTTTTTATGGTAAGTAAGGGTGAAGAAGACAATATGGCGATTATTAAAG
AGTTCATGCGTTTTAAAGTTCACATGGAAGGTAGCGTAAATGGGCATGAATTTGAGATTGAA
GGAGAGGGTGAAGGACGTCCGTACGAGGGTACCCAAACCGCGAAGCTCAAAGTTACAAAA
GGCGGTCCACTTCCATTTGCTTGGGATATTCTTAGCCCGCAATTCATGTATGGTTCAAAAGC

ATATGTGAAACATCCAGCGGACATCCCTGATTATCTGAAGTTATCTTTTCCGGAAGGTTTTAA
ATGGGAGCGTGTTATGAACTTCGAAGACGGCGGGGTCGTGACGGTGACACAAGATTCCTC
GCTGCAAGATGGTGAGTTCATTTACAAAGTGAAGCTCCGCGGTACAAATTTTCCGTCTGATG
GACCTGTCATGCAAAAGAAGACGATGGGTTGGGAAGCTTCAAGCGAGAGAATGTACCCGG
AAGATGGTGCCCTTAAAGGTGAGATTAAACAACGTCTTAAATTGAAAGACGGTGGTCACTAC
GACGCGGAAGTTAAGACTACATACAAAGCAAAGAAGCCAGTCCAATTACCAGGTGCTTATAA
TGTTAACATTAAGTTAGACATCACCAGTCACAATGAGGATTACACGATTGTTGAGCAATATGA
AAGAGCTGAAGGACGTCACTCGACCGGTGGGATGGACGAGCTTTATAAAGGAGGAGGTGG
TTCCGGCGAGAATTTATATTTCCAAGGCGGCGGCGGCGGTTCGGGTGGGGTCGCAGATCT
CATTAAGAAATTTGAATCCATTTCTAAAGAAGAAGGGGGGGGCGGTAGCGGGCATCATCATC
ATCATCATTGA

**TRAP-HiBiT:**

Ribosome-binding site

Cell-penetrating peptide

HiBiT

GGGSG linker

TEVp cleavage site

Dark mCherry (soluble tag)

GSG linker

Lifeact

6xHis tag

AGAAAGTAAGGAGGTTTTTTATGCGCCAGATTAAAATTTGGTTCCAGAACAGAAGAATGAAAT
GGAAGAAGGGCGGTGGTGGATCAGGTGTCTCAGGGTGGCGCCTTTTCAAGAAAATATCTG
GCGGTGGTGGCAGCGGCGAAAACCTTTACTTTCAGGGCGGTGGTGGTGGTTCGGGAGAG
GAAGATAATATGGCCATAATTAAAGAATTTATGCGCTTCAAAGTTCATATGGAAGGGTCTGTAA
ATGGTCATGAATTTGAAATAGAAGGGGAAGGTGAAGGACATCCGTATGAAGGGACTCAAAC
AGCTAAACTCAAAGTAACAAAAGGCGGTCCGCTTCCGTTTGCTTGGGATATTTTATCCCCTC
AATTTATGTATGGATCAAAAGCGTATGTCAAACATCCGGCTGATATACCGGATTATTTGAAACT
TAGCTTTCCAGAAGGTTTTACTTGGGAACGTGTTATGAATTTTGAGGATGGTGGAGTCGTCA
CTGTCACGCAAGATTCTAGTCTTCAAGATGGGCAGTTTATATATAAGGTTAAACTCTTAGGAAT
AAATTTTCCGTCGGATGGGCCAGTTATGCAAAAGAAAACTATGGGATGGGAAGCATCAACAG
AACGCATGTATCCGGAAGATGGTGCTTTGAAAGGGGAAATTAATCAACGCCTTAAATTAAAA
GATGGTGGGCATTATGATGCCGAAGTTAAAACAACATATAAAGCTAAGAAACCAGTACAGCT
GCCTGGGGCGTATAATGTTGATATTAAACTCGATATAACGAGTCATAATGAAGATGGTTCTGG
CGGAGTAGCTGACTTGATTAAGAAGTTCGAGAGTATCAGCAAGGAAGAGGGCGGCGGCTC
CGGACATCATCATCATCATCATTGA

**TRAP-ΔCPP-HiBiT:**

Ribosome-binding site

HiBiT

GGGSG linker

TEVp cleavage site

Dark mCherry (soluble tag)

GSG linker

Lifeact

6xHis tag

AGAAAGTAAGGAGGTTTTTTATGGTTAGCGGTTGGCGCTTATTTAAGAAAATTAGTGGTGGT
GGTGGGTCAGGCGAAAACCTTTACTTTCAGGGCGGCGGCGGTGGCTCTGGAGAAGAAGAT
AATATGGCGATTATTAAAGAATTTATGCGTTTTAAAGTTCATATGGAAGGAAGTGTTAATGGTC
ACGAATTTGAAATTGAAGGGGAAGGTGAGGGGCATCCTTATGAAGGAACGCAAACAGCAAA
ATTAAAAGTTACTAAAGGCGGTCCTCTTCCATTTGCTTGGGATATATTGTCACCGCAATTTATG
TATGGGAGTAAAGCGTATGTAAAACATCCAGCTGATATTCCGGATTATCTCAAACTCTCTTTTC
CTGAAGGATTTACATGGGAACGGGTTATGAATTTTGAAGACGGTGGAGTAGTTACAGTCACA
CAAGACTCATCCCTTCAAGATGGACAGTTTATTTATAAAGTTAAATTGCTTGGAATTAATTTCC
CAAGTGACGGTCCAGTAATGCAAAAGAAAACGATGGGGTGGGAAGCATCTACTGAACGTAT
GTATCCAGAAGATGGTGCACTTAAAGGGGAAATAAATCAACGGCTTAAACTCAAAGATGGTG
GTCATTATGATGCTGAAGTTAAGACGACTTATAAAGCAAAGAAACCTGTACAGTTACCGGGT
GCTTATAATGTAGATATAAAACTCGATATTACATCCCACAATGAAGATGGGTCAGGTGGCGTG
GCCGACCTCATCAAGAAGTTCGAGTCAATCAGCAAGGAAGAGGGTGGCGGCAGCGGCCAT
CATCATCATCATCATTAA

**Supplementary Figures**

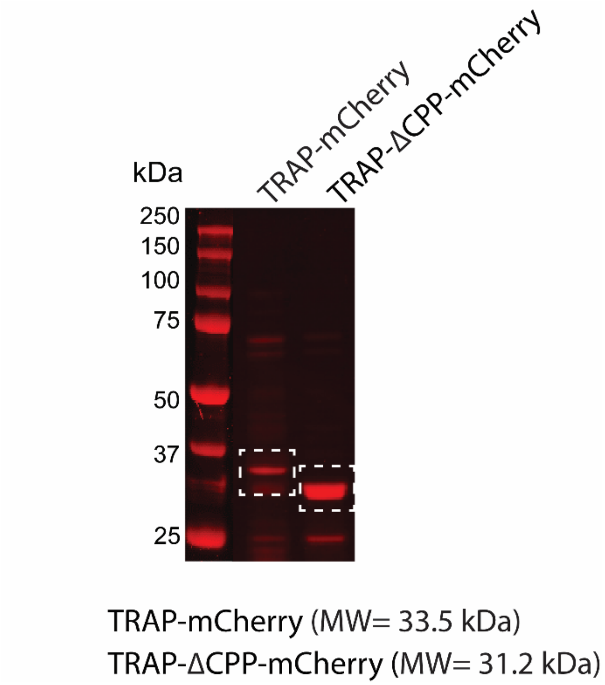


**SI Figure 1.** SDS-PAGE image (by fluorescence gel scanner) showing the purified TRAP-mCherry and TRAP-ΔCPP-mCherry after elution from Ni-NTA resin with 250 mM imidazole. Left lane shows the ladder.

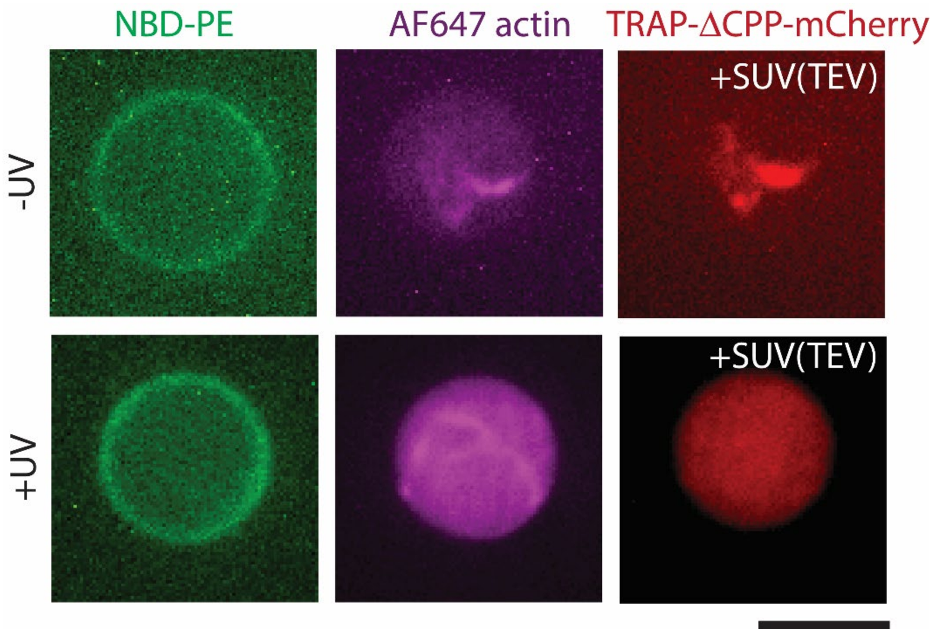


**SI Figure 2.** Representative confocal fluorescence microscopy images of GUVs (green) encapsulating F-actin (magenta), DC(8,9)PC{TEVp} SUVs along with TRAP-ΔCPP-mCherry (red) with or without 10 min exposure to UV light. Scale bar: 10 µm.

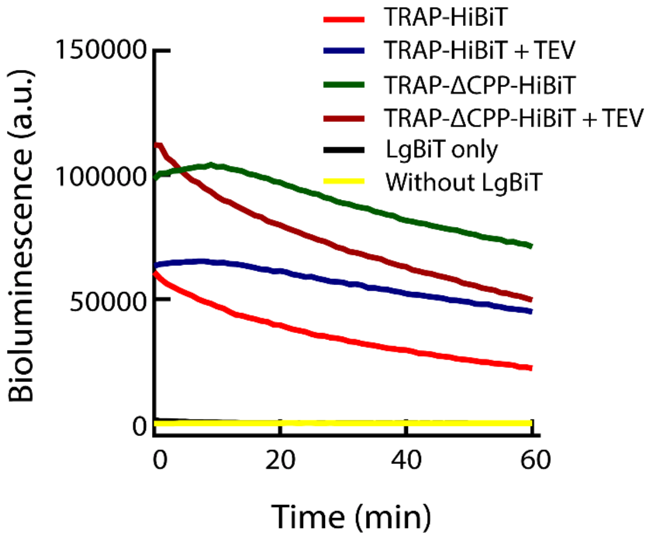


**SI Figure 3.** Time-dependent luminescence measurements of 1 μM TRAP-HiBiT or TRAP-ΔCPP-HiBiT mixed and incubated for 10 min with 1 μM LgBiT in the presence or absence of TEVp. Also shown is the luminescence measurement from TRAP-HiBiT without prior mixing and incubation with LgBiT.

**Supplementary Table**

**Table S1: List of p-values**

| Figure | Condition | Comparison values | p-values |
|---|---|---|---|
| **3-f** | TRAP-ΔCPP-mCherry | 0 min vs 30 min | 0.6631 |
| **3-g** | TRAP-mCherry | 0 min vs 30 min | $2.87\times10^{-7}$ |
| **4-d** | -DC(8,9)PC[TEVp]-UV | TRAP-ΔCPP-mCherry vs TRAP-mCherry | 0.9205 |
| | -DC(8,9)PC[TEVp]+UV | TRAP-ΔCPP-mCherry vs TRAP-mCherry | 0.6791 |
| | +DC(8,9)PC[TEVp]-UV | TRAP-ΔCPP-mCherry vs TRAP-mCherry | 0.0071 |
| | +DC(8,9)PC[TEVp]+UV | TRAP-ΔCPP-mCherry vs TRAP-mCherry | $<1.0\times10^{-12}$ |
| | TRAP-mCherry | -DC(8,9)PC[TEVp]-UV vs -DC(8,9)PC[TEVp]+UV | 0.0030 |
| | TRAP-mCherry | -DC(8,9)PC[TEVp]+UV vs +DC(8,9)PC[TEVp]-UV | 0.9988 |
| | TRAP-mCherry | +DC(8,9)PC[TEVp]-UV vs +DC(8,9)PC[TEVp]+UV | $2.1048\times10^{-8}$ |
| **4-f** | TRAP-ΔCPP-HiBiT | -DC(8,9)PC[TEVp] vs +DC(8,9)PC[TEVp] | 0.2108 |
| | TRAP-HiBiT | -DC(8,9)PC[TEVp] vs +DC(8,9)PC[TEVp] | $<1.0\times10^{-12}$ |
